# Non-random inversion landscapes in prokaryotic genomes are shaped by heterogeneous selection pressures

**DOI:** 10.1101/092056

**Authors:** Jelena Repar, Tobias Warnecke

## Abstract

Inversions are a major contributor to structural genome evolution in prokaryotes. Here, using a novel alignment-based method, we systematically compare 1651 bacterial and 98 archaeal genomes to show that inversion landscapes are frequently biased towards (symmetric) inversions around the origin-terminus axis. However, symmetric inversion bias is not a universal feature of prokaryotic genome evolution but varies considerably across clades. At the extremes, inversion landscapes in Bacillus-Clostridium and Actinobacteria are dominated by symmetric inversions, while there is little or no systematic bias favouring symmetric rearrangements in archaea with a single origin of replication. Within clades, we find strong but clade-specific relationships between symmetric inversion bias and different features of adaptive genome architecture, including the distance of essential genes to the origin of replication and the preferential localization of genes on the leading strand. We suggest that heterogeneous selection pressures have converged to produce similar patterns of structural genome evolution across prokaryotes.

---

In both eukaryotes and prokaryotes, genome architecture and its evolution are frequently non-random^1,2^. A fundamental question in this regard is whether non-random genome organization is brought about by biased mutational processes, for example relating to the dynamics of recombination, or by selection. In prokaryotes, many aspects of non-random genome organization have been plausibly attributed to the latter. This includes the clustering of functionally related genes into operons and the enrichment of essential genes on the leading strand of replication, where they avoid head-on collisions between active DNA and RNA polymerases^3–5^. In addition, highly expressed genes (rRNA, tRNA, ribosomal protein genes) are commonly found near the origin of replication *(ori)* in both bacteria and archaea^6–8^, consistent with selection for elevated dosage: sequences near *ori* replicate earlier and are therefore transiently present in higher copy number, a phenomenon exacerbated in fast-growing organisms where multiple rounds of replication can be initiated concurrently.

For other aspects of prokaryotic genome evolution, it has been more difficult to pin down whether non-random genome structure is brought about by selection, biased mutational processes or a combination of the two. This notably includes the incidence pattern of large-scale inversions, which constitute a major source of structural diversity in prokaryotic genomes^9,10^. Curiously, inversions in prokaryotes appear to be predominantly symmetric (Figure 1A); that is, their end points are approximately equidistant from the origin of replication, generating conspicuous X patterns (Figure 1B) in whole-genome alignments^11–14^. Inversions symmetric to the origin-terminus *(ori-ter)* axis were initially observed in a few closely related bacterial genomes (including pairs of Chlamydia, Mycobacterium, and Helicobacter *spp*.^11–14^) and have subsequently been highlighted in multiple other genome comparisons, notably involving γ-proteobacteria (Yersinia^15^, Blochmannia^16^, Buchnera^17,18^, Pseudomonas^19^) and Bacilli (Lactobacillus^20^, Bacillus^19^, Streptococcus^21^), but also the single-origin archaeon Pyrococcus^22^. These observations have given rise to the notion that biased inversion landscapes are a prevalent feature of prokaryotic genome evolution.

**Figure 1.**
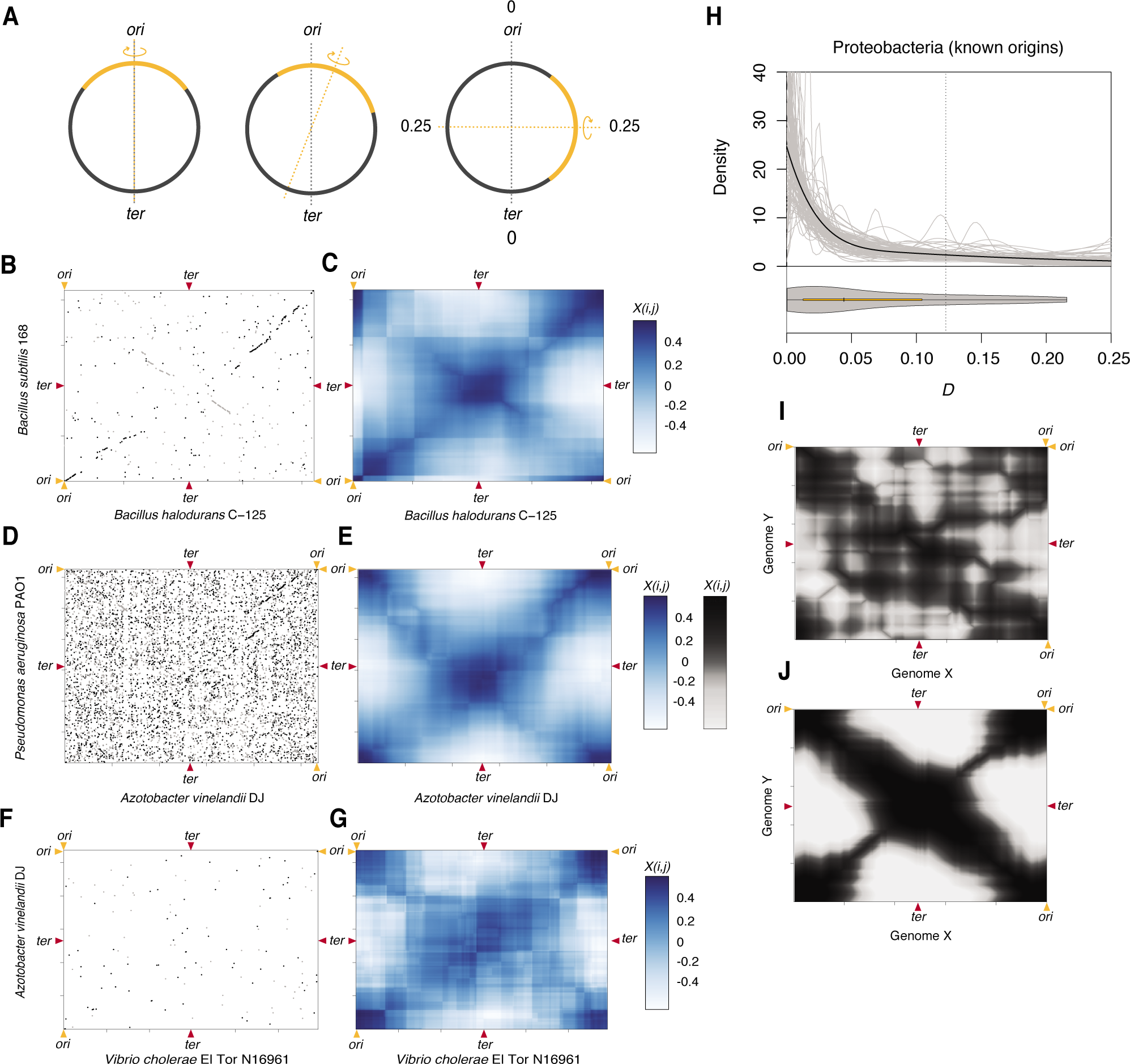
Detecting symmetric inversion bias. (A) Inversions in circular prokaryotic genomes can occur with varying degrees of symmetry in relation to the *ori-ter* axis (grey dotted line). The furthest away the inversion axis (orange dotted line) can be from the *ori-ter* axis is a quarter of the genome (0.25). (B, D, F) Symmetric inversions cause X-shaped patterns in pairwise MUMmer alignments, which can be hard to discern (D, F) but are revealed as symmetry hot and cold spots when displaying *X*_*i,j*_ (C, E, G). (H) Distribution of *D* scores for Proteobacteria with experimentally determined *ori* positions. The black line is the density fit to the whole dataset, the grey lines are density fits to random 40% jack-knifed samples to explore outsize influence of individual genome pairs. (I, J) *X*_*i,j*_ heat maps representing a case of simulated genome divergence by symmetric and quasi-symmetric (I) and random (J) inversions.

Both non-random mutational processes and selection have been suggested as potential drivers of biased inversion landscapes. Regarding the former, it has been proposed that symmetric inversions might be favoured by the layout of bacterial replication. As sister replisomes progress at approximately the same speed after setting out together from the origin of replication, homologous recombination across replichores would lead to symmetric or quasi-symmetric inversions if single-stranded DNA, present in the wake of one of the sister replisomes, is used as a template for illegitimate recombination following the generation of a double strand break in the vicinity of the other replisome^12,23^.

Alternatively, the genesis of inversions is approximately random but symmetric inversions are more likely to survive purifying selection because they are, on average, less disruptive to adaptive genome architecture^11,12,24^. Notably, although a potentially large number of loci are translocated to the opposite replichore, they will retain their original leading/lagging strand orientation. This might be important not only to avoid replication-transcription conflicts but also for binding motifs that function in a polarized fashion, such as FtsK-orienting-polar-sequences, which facilitate FtsK translocation towards the terminus. In contrast, inversions *within* the same replichore inevitably result in leading/lagging strand switches. Symmetric inversions also do not alter the distance of a particular genomic element to *ori* or *ter*, thus avoiding potentially deleterious changes in gene dosage and the displacement of motifs whose function is contingent on their proximity to either *ori* or *ter* (e.g. DnaA boxes, parS motifs)^25,26^. Finally, symmetric inversions do not alter relative replichore length, which is important in light of experimental evidence from *E. coli* that replichore size imbalance of more than 10% is deleterious^27^.

Here, to elucidate the relative importance of selection and mutation bias in the evolution of inversion landscapes across prokaryotes, we first take a step back and ask: are symmetric inversions really a universal feature of prokaryotic genome evolution? The tally of individual cases in the literature seems striking, yet symmetric inversions have - with few notable exceptions^11.15.28^ - been diagnosed rather casually by picking out apparent X patterns from pairwise whole genome alignments by eye. This is problematic, not least because there might be extensive reporting bias, i.e. obvious symmetric inversions are highlighted in the literature, whereas subtle biases are missed and random inversion patterns rarely mentioned^29^.

To address this issue, we developed an alignment-based approach to systematically assess inversion symmetry between pairs of prokaryotic genomes. Applying this approach to 1651 bacterial and 98 archaeal genomes, we demonstrate that there is substantial heterogeneity in the prevalence of symmetric inversions across prokaryotic clades. While inversion landscapes in some phyla, for example Bacillus-Clostridium, are dominated by symmetric inversions, the propensity to invert around the *ori-ter* axis is much less pronounced in other clades, including Bacteroidetes and Euryarchaeota. We go on to show that putatively adaptive features of genome architecture linked to the *ori-ter* axis, such as the fraction of genes encoded on the leading strand and the average distance of rRNA genes to the origin of replication, are predictive of symmetric inversion bias but in clade-specific fashion. For example, enrichment of highly expressed informational genes near *ori* is predictive of symmetric inversion bias in Proteobacteria, but not in Actinobacteria, where we observe a strong correlation with relative nucleotide motif abundance on the leading versus lagging strand instead. We suggest that heterogeneous selection pressures have converged to produce similar patterns of genome evolution. Dissecting inversion patterns as a function of known replication dynamics in different organisms, we also suggest that selection might frequently act on top of and reinforce pre-existing mutational biases.

## Results and Discussion

To assess inversion symmetry across a large number of genomes, we developed a simple geometric approach for pairwise genome comparisons that is fast, easily scalable, and applicable across different phylogenetic distances. Starting from MUMmer^30^ alignments of two genomes, like the one shown in Figure 1B, we make use of the fact that, if there is a single dominant axis around which inversions have occurred between two genomes, homologous sequence blocks will be located on one of the two diagonals that pass through that axis, generating the familiar X pattern. We can test how well the distribution of homologous sequences in the two genomes conforms to this single-symmetry-axis model by considering, in a strand-specific fashion, the residual deviation of homologous sequence blocks from the expected X pattern. Briefly, rather than define a likely symmetry axis (e.g. *ori-ter),* we systematically survey the MUMmer alignment landscape at a resolution of 10kb and, for each point *i,j,* register the strength of X-type symmetry (X_i,j_) by considering the position of homologous blocks with respect to an imaginary X centred at *i,j.* More formally, X-type symmetry is defined as:

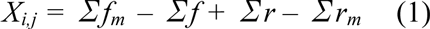

where *f* and *r* are residual deviations from the forward arm of the imaginary X of homologous blocks in forward and reverse orientation, respectively; and *f*_*m*_ and *r*_*m*_ are residual deviations of the same homologous blocks mirrored around the vertical axis going through *i,j.* The logic here is the following: An inversion around *i,j* moves a given sequence onto the other arm of the X. Mirroring should reverse this step, in effect restoring collinearity in the genome alignment, which will minimize *r*_*m*_ or *f*_*m*_ and therefore lead to a large X_i,j_. This approach effectively converts the MUMmer alignment into a matrix of symmetry hot and cold spots (Figure 1C, E, G). Visualizing such matrices frequently reveals strong X-type symmetry (henceforth simply referred to as symmetry), including in genomes that are substantially diverged so that homologous sequences are sparse and in genomes that exhibit high rates of rearrangement and where symmetry would have likely escaped visual detection (Figure 1D–G). When simulating inversion events between two dummy genomes, very similar patterns are evident when rearrangements are restricted to symmetric or quasi-symmetric inversions (Figure 1J) whereas the tell-tale secondary diagonal is not present when no symmetry constraints are imposed and inversions can occur at random throughout the genome (Figure 1I, see Methods for details).

Up to this point we have identified points of inversion symmetry in an unbiased fashion without regard to genomic landmarks such as *ori* or *ter*. To test empirically whether inversion symmetry around the *ori-ter* axis is prevalent across prokaryotic genomes, we calculated the distance (D) between the point of maximal symmetry [max(X_i,j_)] in a given MUMmer alignment and *ori* or *ter* (whichever is closer). First, we considered a small set (N=19) of proteobacterial genomes with experimentally determined origins (supplementary Table 1). Making all pairwise comparisons that satisfy a lenient set of minimum homology criteria (see Methods) we find a clear departure from the null model of random inversions. *D* is strongly shifted towards smaller values, indicative of inversions largely happening around the *ori-ter* axis (Figure 1H).

### Variable prevalence of symmetric inversions across prokaryotic phyla

The location of *ori* can be robustly predicted in many prokaryotic genomes based on a set of hallmark features, including the location of DnaA boxes and strand biases in nucleotide composition that change sign around *ori*, reflecting divergent mutational biases associated with leading/lagging strand replication. We therefore extended our analysis to include genomes with computationally predicted origins (see Methods). In total, we inferred symmetry scores and calculated *D* values for 528396 (2645) pairwise comparisons between 1651 bacterial (98 archaeal) genomes with predicted *ori* coordinates (supplementary Table 2). We then considered the distribution of *D* values aggregated at the phylum level. We find strong symmetric inversion bias in multiple bacterial phyla, including those that have shaped our understanding of replication-associated genome architecture - Proteobacteria and Bacillus-Clostridium (Figure 2) - in line with previous observations from individual genome pairs and a more systematic comparison of recently diverged genomes^28^. However, *ori*-ter biased inversion landscapes are not universal and the degree of bias varies considerably between clades. For example, the rearrangement landscapes of Bacteroidetes-Chlorobi-Fibrobacteres (BCF) and Tenericutes show much weaker bias towards symmetric inversions than other clades, and we detect no significant biases in archaea with a single origin of replication (Figure 2). Heterogeneity in symmetric inversion bias is also evident at lower taxonomic levels, as illustrated for Proteobacteria in Figure 2.

**Figure 2.**
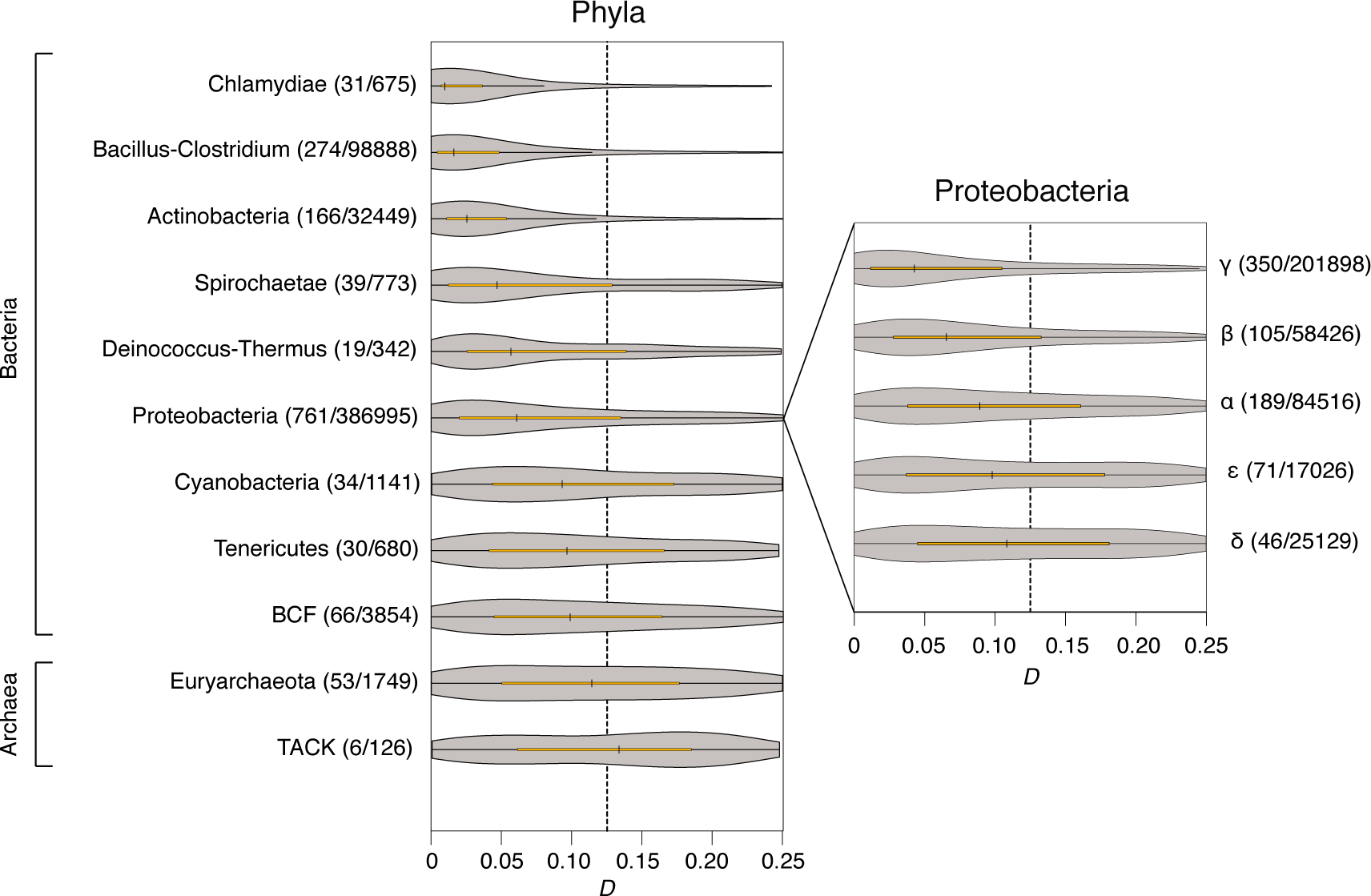
Distribution of *D* scores in different prokaryotic clades. The number of focal species and the number of eligible pairwise comparisons within each phylum or class is given in parentheses. Note that, while at least one of the two paired genome belongs to the indicated clade, its partner is chosen based on the presence of a minimum number of homologous blocks (see Methods) and might therefore in some instances belong to a different clade.

*D* values support symmetric inversion bias across a remarkably broad range of divergence levels. However, weaker *ori-ter* symmetry signals are generally observed for very closely and distantly related genomes, as exemplified in Figure 3A by the phylum Bacillus-Clostridium. The variability in symmetry scores over evolutionary time might be the consequence of limited rearrangement signal at short evolutionary distance and lower signal-to-noise ratio when divergence levels are high. To rule out that heterogeneity in symmetric inversion bias between clades is the consequence of differential sampling of divergence levels, we subsampled from within each individual phylum to match a common distribution of 16S rRNA distances (see Methods). Taking phylogenetic distances into account suggests that we might overestimate the extent of symmetric inversion bias in Chlamydiae. However, globally the results demonstrate that differences in *D* score distributions between clades are robust (Figure 3B).

**Figure 3.**
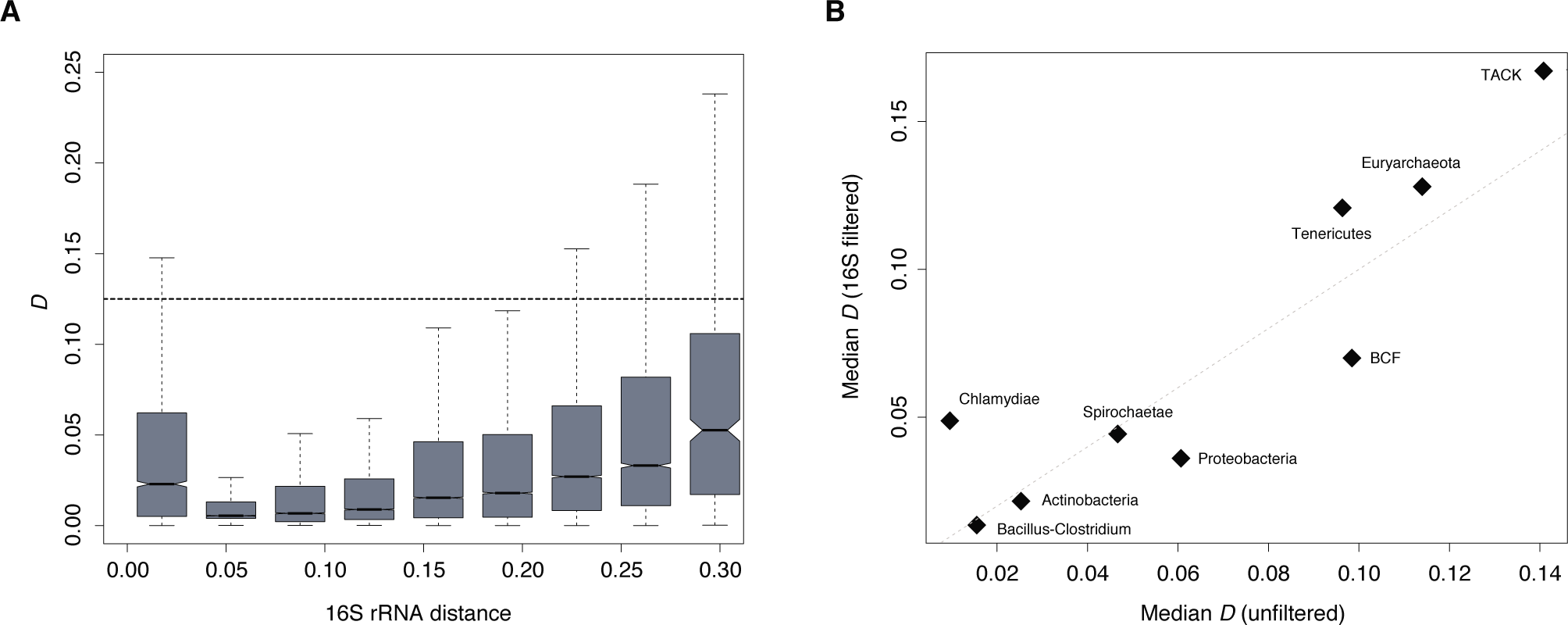
(A) *D* as a function of phylogenetic distance between the aligned genome pairs in the Bacillus-Clostridium phylum. (B) Median symmetry scores (D) for each phylum considering either all pairwise genome comparisons (unfiltered) or a subset of pairwise comparisons sampled to match a common underlying distribution of 16S divergence levels (16S filtered) as described in Methods (ρ=0.8, P=0.01).

### Selection is a prominent driver of symmetric inversion landscapes

To test whether biased inversion landscapes are likely the result of natural selection to preserve adaptive genome architecture, we considered four features of genome organization linked to the *ori-ter* axis and therefore potentially predictive of symmetric inversion bias: the average distance of rRNA genes to *ori*; the fraction of genes located on the leading strand of replication; the enrichment of highly expressed translational genes (COG J) near *ori*; and the biased distribution of nucleotide motifs - which may be involved in DNA replication, repair or segregation - on either the leading or lagging strand. Regarding the latter, we followed prior work^26^ and confined analysis to octameric nucleotide motifs. We then correlated these features with the median *D* values of the focal genome, computed across all pairwise comparisons in which it participates. Focusing on phyla represented by at least 30 taxa, we find strong correlations with all four features in the expected direction for the phyla Bacillus-Clostridium and Proteobacteria: Genomes with higher symmetric inversion bias (lower median D) have stronger enrichment of COG J genes near the origin of replication (more negative Z score), have rRNA genes encoded at a smaller average distance to *ori*; have a greater fraction of genes encoded on the leading strand of replication and exhibit stronger imbalances of nucleotide motifs (Figure 4). Surprisingly, Actinobacteria - a phylum with a stronger symmetric inversion bias than Proteobacteria - show no intra-phylum correlation between *D* and either COG J gene enrichment or rDNA distance or biased gene orientation (Figure 4). However, there is a strong relationship in this clade between the degree of octamer strand bias and median *D*. A similar pattern is evident for the BCF phylum, where, in addition, we observe a relationship with rDNA distance. In contrast, biased distribution of genes on the leading strand is the only feature predictive of symmetric inversion bias in Tenericutes. Overall, these results suggest that symmetric inversion bias is promoted by different principal selection pressures in different prokaryotic clades.

**Figure 4.**
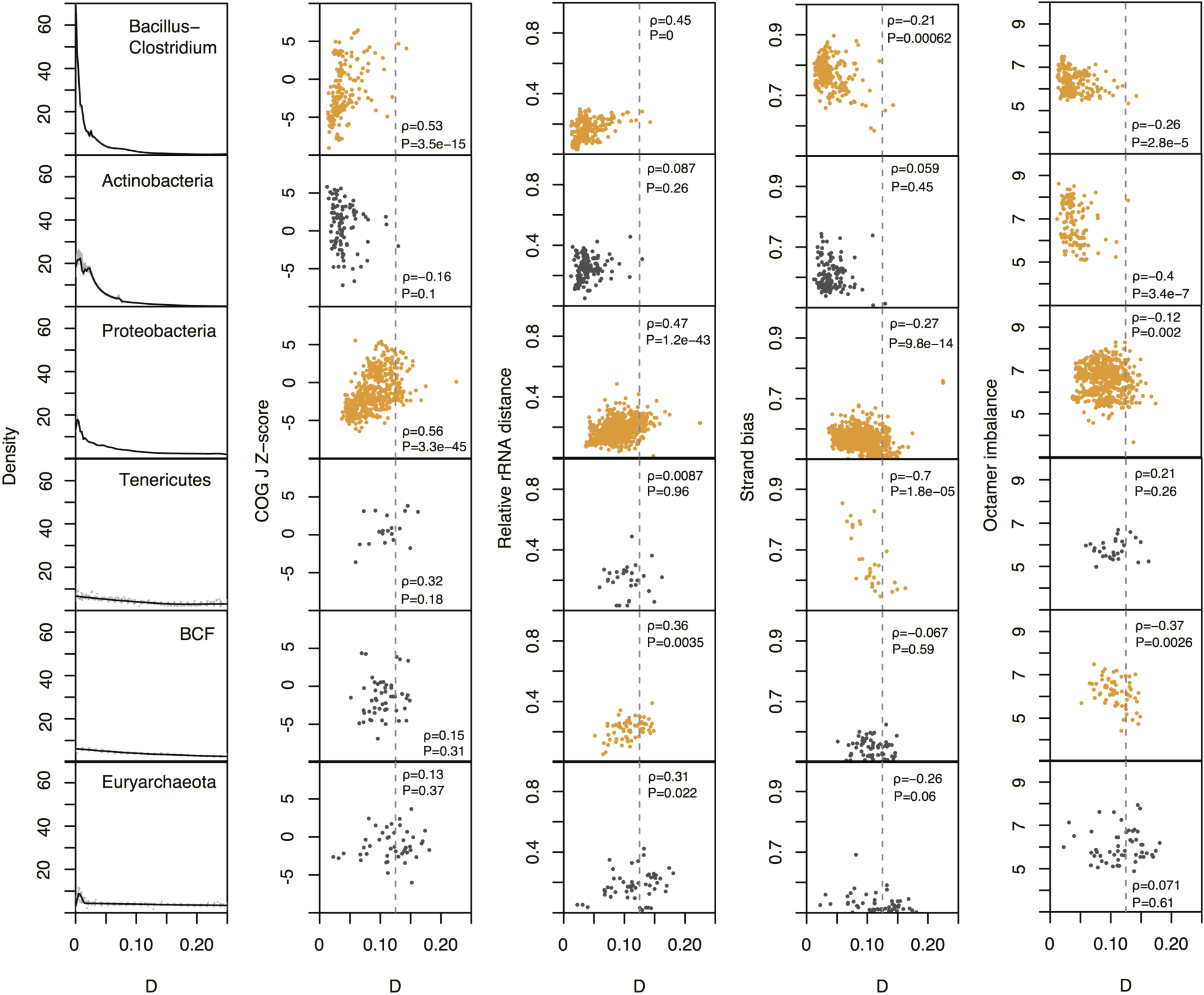
Relationship between *D* and different features of *ori-ter* related adaptive genome architecture. Correlations below a P value threshold of 0.005 are highlighted in orange. See main text for a description of the covariates.

### Sister replisome proximity predicts symmetric inversion bias at short evolutionary distances

Despite inter-clade heterogeneity, the analyses above argue that selection is pervasively implicated in shaping inversion landscapes across prokaryotic evolution. But does selection operate on an initially random population of inversions or does it act to reinforce pre-existing mutational biases? In other words, is the mutational raw material already biased in favour of symmetric inversions? The sister replisome hypothesis of symmetric inversions assumes that physical proximity between sister replication forks facilitates illegitimate recombination and predicts that symmetric inversions should be more likely in organism with higher levels of sister fork co-localization.

To test this hypothesis, we considered variability in physical proximity of sister replication forks during the replication cycle, which has been examined by fluorescence microscopy in a range of different bacteria. In some species, sister replisomes associate tightly throughout the replication cycle, notably in *Bacillus subtilis*^31^ (Firmicutes) and *Pseudomonas aeruginosa*^32^(γ-proteobacteria). where they co-localize at mid-cell, and in *Caulobacter crescentus*^33^ (α- proteobacteria) and *Helicobacter pylori*^34^ (ε-proteobacteria), where sister forks remain in close physical proximity when they migrate together from the cell pole to a mid-cell position. Sister replisomes are not physically tethered to each other (they move independently) but remain close. In *Mycobacterium smegmatis*^35^ (Actinobacteria) and *Myxococcus xanthus*^36^ (δ-proteobacteria) sister replisomes sporadically drift far enough apart to allow detection of independent fluorescence foci but stay within a certain distance of each other^35,37^. Finally, in *Escherichia coli,* replisomes have been reported to localize to different cell poles after initiation of replication at mid-cell^38^. However, it remains unclear to what extent independent foci in the same cell might represent a further round of replication initiation and more extensive tracking of individual sister replisomes over time is required to understand co-localization throughout the cell cycle. With these caveats in mind, we find that *D* scores correlate positively with the fraction of observations in a given replisome tracking experiment that detected two foci rather than a single focus (Figure 5, ρ=0.9, P=0.08, Spearman’s correlation, supplementary Table 3).

**Figure 5.**
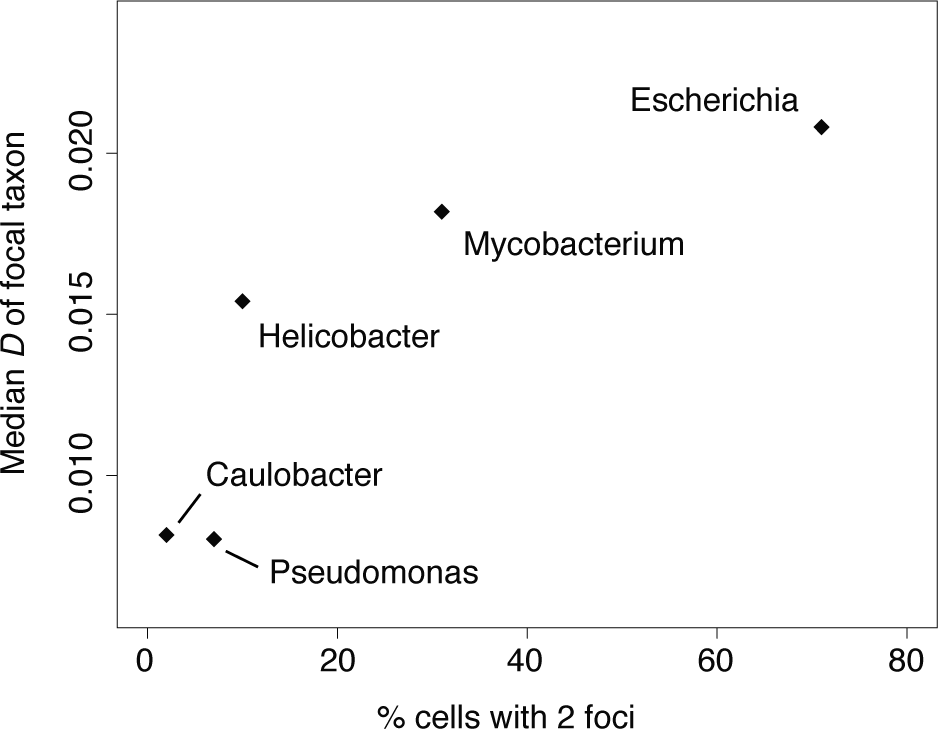
Relationship between *D* and the fraction of cells with two fluorescent replisome foci. *D* scores were calculated for closely related taxa (see main text). The fraction of cells showing two fluorescent foci based on replisome tracking experiments for *Pseudomonas aeruginosa*^32^, *Caulobacter crescentus*^33^, *Helicobacter pylori*^34^, Mycobacterium smegmatis^35^ *and* Escherichia coli^38^.

That is, organisms in which putative sister replisomes spend more time apart and are thus detectable as independent foci, have a lesser tendency for symmetric inversions. In computing *D* score here, we focus on inversions between closely (strain level) related genomes (16S rRNA divergence <0.01) because a) it increases the chance that we are dealing with mutational events that have not yet been eliminated by selection and b) we can reasonably assume that co-localization dynamics in the compared taxa are the same whereas this need not be the case between more distantly related taxa, as illustrated by the different co-localization patterns of *P. aeruginosa* and *E. coli* (both γ-proteobacteria). These findings hint at the possibility that selection operates on pre-existing mutational biases to reinforce and maintain biased inversion landscapes. However, we note that the small number of species for which we have data on replisome dynamics limits our ability to draw firm conclusions in this regard.

### Symmetric inversions are prevalent around termini in archaea with multiple origins of replication

Importantly, however, sister replisome co-localization is not necessary for symmetric inversion bias to be observed. We can demonstrate this explicitly by considering inversion dynamics in archaea with multiple origins of replication. In single-origin prokaryotes symmetry around the origin is inextricably tied to symmetry around the terminus. In multi-origin organism, on the other hand, it is possible in principle to observe symmetric inversions around a given terminus but not a neighbouring origin, and vice versa. If sister replisomes travelling together from the origin provide the main mutational impetus for symmetric inversions, symmetry should be evident around origins, but not around termini of replication. Selection on the other hand, might favour symmetric inversions both around the origin and the terminus, for example to maintain a given distance to the terminus for motifs involved in DNA segregation. We therefore considered inversions around the origins and termini of replication in Sulfolobus *spp.* (3 origins) and Halobacteria (3-4 origins), the two clades where multiple complete genomes are available for comparison. In both *Haloferax volcanii* and *Sulfolobus acidocaldarius* - model representatives of these clades - origins of replication fire synchronously and evidence from *Sulfolobus acidocaldarius* suggests that sister replisomes remain associated during replication^39^. At the same time, despite the resemblance to eukaryotic, multi-ori mode of chromosome replication, selection has shaped the organization of Sulfolobus genomes in a bacteria-like fashion, with essential and highly expressed genes enriched around the origins of replication^7^. We divided 15 Sulfolobus and 14 halobacterial chromosomes into *ori*- or *ter*-centered territories, and paired up orthologous territories from different taxa to investigate the presence of the X-type symmetry in each territory (see Methods).

Symmetric inversion bias is evident around individual origins of replication but, strikingly, also around individual termini in both clades (Figure 6). As symmetric inversions around replication termini cannot have originated from co-travelling sister replisomes, we conclude that either selection or an alternative mutational bias must account for the non-random inversion landscape.

**Figure 6.**
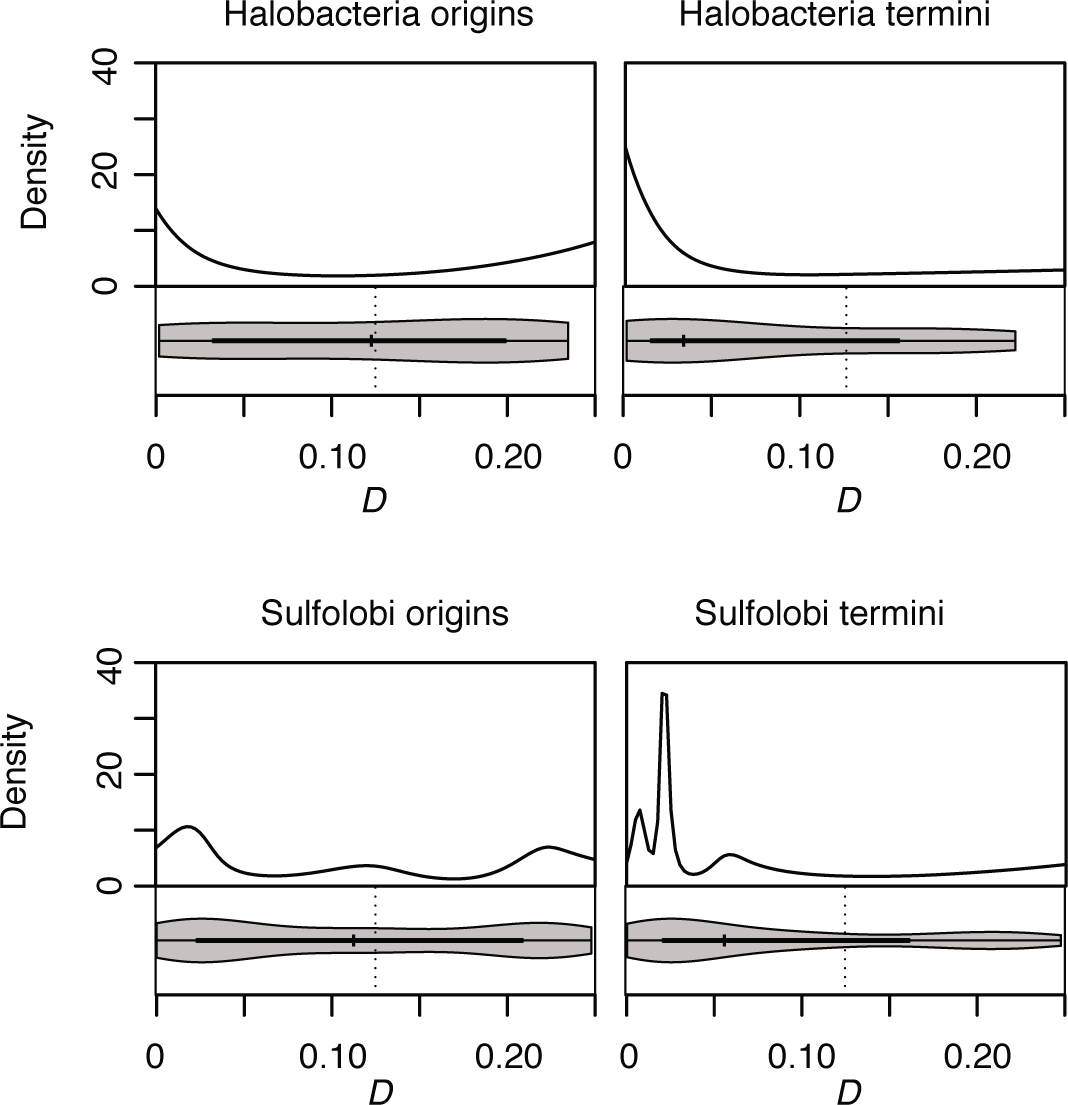
Distribution of *D* scores for *ori-* and fer-centered territories in Sulfolobus and Halobacteria.

Regarding alternative mutational biases, one intriguing possibility is that 3D chromosome topology either during or outside of replication affects inversion landscapes. Indeed, it has been suggested recently^28^ that chromosomal arrangements within the 3D cell space might promote symmetric inversions. In most prokaryotes where chromosome conformation has been probed globally, including *C. crescentus*^40–41^*, B. subtilis*^42^ and *Vibrio cholerae*^43^, chromosome topology follows a longitudinal organization, the right and left replichore lying parallel to each other in the cell. If homology search following a double strand break were biased towards regions of the genome that are close in space, a longitudinal organization would favour the emergence of symmetric inversions. Consistent with this hypothesis, Khedkar and Seshasayee (2016) find that regions of high contact density in *C. crescentus* are enriched for inversion breakpoints. This potential mechanistic bias deserves further scrutiny in light of evidence that the 3D arrangement of chromosomes inside the cell is predictive of translocation probabilities in eukaryotes^44^.

While we cannot currently quantify the role different mutational biases play in shaping symmetric inversion bias, our results strongly suggest that selection is the ultimate sculptor of biased inversion landscapes across prokaryotic evolution and that heterogeneous selection pressures can converge to produce similar patterns of genome structure evolution in different prokaryotic clades.

## Methods

### Data acquisition and processing

Genome sequences of 2784 and 2773 completely sequenced prokaryotes were downloaded from Refseq (ftp.ncbi.nlm.nih.gov\genomes\Bacteria, accessed 5 Oct 2014) and GenBank (ftp://ftp.ncbi.nlm.nih.gov/genomes/archive/old_refseq/Bacteria/, accessed 12 Mar 2015), respectively, along with corresponding annotations. Where multiple genome elements were present for a given taxon (secondary chromosomes, plasmids), we only considered the largest chromosome.

We assembled a dataset of Proteobacteria with experimentally determined *ori* position from the literature (supplementary Table 1). Where *oris* are defined by two flanking genes rather than a precise genome coordinate, the centre position between the two genes was assigned as *ori. In silico* predicted *ori* positions were obtained from the DoriC database (http://tubic.tju.edu.cn/doric/), which integrates information about multiple genome features (nucleotide skews, DnaA box distribution, genes adjacent to candidate *oris)* to predict *ori* positions^45^. Where necessary, the NCBI Genome Browser was used to map Refseq genome element identifiers from DoriC to GenBank identifiers. Only genomes with assemblies used to build the DoriC database were included in the analysis. In some instances, DoriC annotates multiple origin locations for a single bacterial chromosome. These cases might constitute genuine multi-partite origins - as observed, for example, in *H. pylori*^46^ - artefacts of feature-based origin calling or represent plasmid integration events. Following manual inspection, all putative origin locations separated by 3500bp or less were considered to be a single origin with multiple parts, and the centre of the region containing multiple parts was taken as *ori.* Bacterial chromosomes with multiple *ori* locations separated by >3500bp were excluded. We defined the terminus as the position half a genome length away from the origin. While not every terminus region in bacteria is located at precisely this position, there is evidence for strong long-term selection to maintain equal replichore size, notably from analyses of nucleotide skews, which suggest that replichore imbalance rarely exceeds a 60:40 split^26^.

### Assessing symmetric inversion bias

To establish whether there are biases for symmetric versus asymmetric inversions in different prokaryotic clades, we first aligned all possible intra-phylum genome pairs using the 'mummer' program from the MUMmer package^30^ to detect maximal unique matches (MUMs) between genome pairs. The length of MUMs varies from a pre-set threshold to the length of the longest exact and unique sequence match detected in both genomes. A threshold of 20bp for the minimal MUM length was chosen based on the recommendations of the authors of the MUMmer package as sufficiently large to avoid spurious matches, and sufficiently small to detect a number of matches in the phylogenetically distant genome pairs. Only genome alignments with at least 40 MUMs in one direction and 20 MUMs in the other direction were retained for analysis.

We then considered the residual deviation of individual MUMs from the forward or reverse arm of an imaginary X spanning the alignment plot, as described in the main text, scanning the alignment space at a resolution of 10kb, and assessing X-type symmetry (X_i,j_) for each point *i,j* according to equation (1). To allow comparisons across genome pairs, each *X*_*i,j*_ value was normalized by the total number of MUMs detected, and residuals were weighed by the ratio of MUM length to minimal MUM length (20bp) so as to place greater confidence in longer homologous blocks.

### Phylogenetic controls

We obtained 16S rRNA alignments from the SILVA database^47^ (v. 119.1) and mapped GenBank identifiers to DoriC Refseq genomes using the NCBI Genome Browser. In case of multiple 16S rRNA sequences for the same taxon, a single sequence (belonging to the largest genome element) was chosen at random. Two bacteria from the dataset with experimentally determined origins of replication were not found in the SILVA alignments *(Rickettsia prowazekii* and *Vibrio harveyi).* For these bacteria, the first 16S rRNA sequence in the GenBank file of the largest genome element was added to the alignment using the SINA aligner available on the SILVA website (https://www.arb-silva.de/). Based on the full 16S rRNA alignment, we calculate pairwise phylogenetic distances between taxa using the 'dnadist' program from the Phylip package (v. 3.695) with default settings. We used NCBI Taxonomy (https://www.ncbi.nlm.nih.gov/taxonomy) to divide species into phylogenetic clades of interest (phyla and classes). To render X-type symmetry values comparable between clades, we randomly subsampled from genome pairs within each clade of interest to match a common template of phylogenetic distances. As the distance template, we used the distribution of phylogenetic distances within the Thaumarchaeota-Aigarchaeota-Crenarchaeota-Korarchaeota (TACK) superphylum, a clade with a relatively small number of genome pairs.

### Covariance between symmetric inversion bias and genome architecture

We assessed four putatively adaptive features of genome architecture to understand whether symmetric inversion bias might be driven by selection. First, we determined the average distance of rRNA genes to the origin of replication (calculated as a percentile of replichore size). Second, we computed the enrichment of protein-coding genes belonging to the COG class J (Translation, ribosomal structure, biogenesis) near *ori* as a Z-score, comparing the average distance of COG J genes to the average distance of all COG-annotated genes. Lower Z-scores indicate a stronger enrichment of COG J genes near *ori.* COG J genes are often highly expressed and essential and therefore taken to represent a class of genes previously identified as enriched near *ori* in some fast-growing bacteria^4^. Third, we determined the relative enrichment of genes on the leading versus lagging strand as an indicator of replication-transcription conflicts. The strand with more genes was considered to be the leading strand. Finally, we considered the relative enrichment of nucleotide motifs on the leading versus lagging strand. Following Hendrickson and Lawrence^26^, we focussed on octamers as informative motifs, identifying the most strand-biased octamer in each genome by computing the difference in abundance between the two replichores. All octamers with a non-AGTC base were excluded from the analysis.

### Ori-and ter-centred territories in archaea with multiple origins of replication

Genomes of multi-ori archaea were separated into either ori-centered territories (with neighbouring *ters* as boundaries of territories) or ter-centered territories (with neighbouring *oris* as boundaries). *Ori* territories were trimmed so that both replichores were of the same length (which is not the case if origins are not equidistant). Homologous territories were defined as those carrying the most MUM hits to each other when all territory pairs between two species are compared, with an additional constraint of maximal difference in length of 10 kb. *D* was measured as the distance of max(*X*_*i,j*_) to the central point of the territory or to the counterpart of the central point half a territory away.

## Acknowledgments

TW is the recipient of intramural funding from the UK Medical Research Council (MRC) and an Imperial College Junior Research Fellowship.

## Author contributions

J.R. carried out all analyses. J.R. and T.W. conceived the study, interpreted the data and wrote the manuscript.

